# Tracking claim changes from preprint to publication across 72,644 biomedical studies using large language models

**DOI:** 10.64898/2026.06.30.735556

**Authors:** Hao Yin, Woosung Ahn, Patrycja M. Forster, Ruslan Rust

## Abstract

Preprints disseminate biomedical research before peer review, but how much those scientific claims change before publication in journals remains uncertain. Here, we matched 72,644 bioRxiv preprints to their peer-reviewed publications by DOI and used a large language model (Claude Sonnet 4.6) to extract one primary and up to two secondary claims from their abstracts. Content change (unchanged, minor, major/substantial) and hedging shift (more cautious, more confident, unchanged) were compared. Four raters independently labelled a 550-pair subset; model–rater agreement (quadratic-weighted Cohen’s κ = 0.69) approached agreement among the raters themselves (κ = 0.76). The primary claims were unchanged in 39.8% of abstract pairs, minorly revised in 50.0%, and substantially revised in 10.2%. Claims became more cautious twice as often as more confident (8.4% vs 4.2%). Substantial revisions correlated with longer preprint-to-publication interval. Therefore, among bioRxiv preprints that are later published, central abstract claims remain stable through the preprint-to-publication transition.

## Introduction

Preprints have changed the timing and avenues of scientific communication. BioRxiv alone has hosted more than a quarter of a million manuscripts. Early analyses showed that 65-70% of bioRxiv preprints posted between 2013 and 2016 were eventually published in peer-reviewed journals^1,2^. A 2025 openRxiv report estimated that approximately 80% of preprints posted to bioRxiv or medRxiv were subsequently published^3^. Preprint use in the biological sciences began to accelerate around 2016, correlating with the establishment of ASAPbio and the Zika epidemic^4^. The COVID-19 pandemic further broadened preprint adoption, particularly in clinical research ^5,6^. Biomedical findings often become publicly available months or years before they appear in journals, allowing researchers and policy makers to act on scientific claims before formal evaluation, usually peer review, is complete. This acceleration also raises uncertainty about how reliable preprint claims are.

Existing studies have reported mixed results. Abstract conclusions change in only 7.2% of non-COVID-19 preprints and in 17.2% of COVID-19 preprints^7^. In another study, results changed importantly between the preprint and its updated or published version in 21% of studies^8^. Yet, effect estimates are largely consistent between preprints and publications^9^ and reporting quality improves only modestly after peer review^10^. Prior studies that examine claims directly are small or COVID-19 specific, and synthesis across them is limited by heterogeneous measures^11–14^. Work that operates at scale instead often measures textual similarity rather than the content of the scientific claims^15,16^, which cannot fully reveal whether a claim was strengthened, weakened, or even overturned. Despite these lines of valuable data, it remains unresolved whether and how the central claims of biomedical research change during the transition from preprints to publications.

Here, we assess this transition across 72,644 bioRxiv preprints posted between 2018 and 2025 that could be matched by DOI to its peer-reviewed version. Using a large language model (LLM, Claude Sonnet 4.6) and validation against human expert raters, we parsed each abstract pair into one primary and up to two secondary claims and classified content, certainty and claim types, which enabled quantitative analysis of the extent and structure of claim-level changes.

We find that, among the preprints that subsequently reached journal publication, approximately 90% of central abstract claims were unchanged or underwent only minor revision, whereas 10% underwent substantial/major revision. Thus, within this publication-selected subset, preprints generally provided a stable early representation of the scientific claims that subsequently appeared in journal articles.

## Results

### Study design and LLM validation

We assembled every bioRxiv preprint posted between 2018 and 2025 that we could match by DOI to its peer-reviewed publication, yielding in total 72,644 preprint-publication pairs (**Supplementary Fig. 1**). For each pair, a large language model (Claude Sonnet 4.6) parsed the preprint and published abstracts into one primary and up to two secondary claims and classified the pair for content change (unchanged, minor, or substantial/major) and for hedging shift (more cautious, more confident, or unchanged). The primary claims were classified by the LLM into exactly one of six mutually exclusive types: mechanistic (with a causal mechanism or mediator), associative (with a non-causal relationship), descriptive (with an observation or characterization), methodological (with the performance of a method), therapeutic (with the effect of an intervention or therapeutic target), or null result (with the absence of an association or effect as the central claim). We validated the model against four independent human raters on a stratified subsample of 550 pairs; this subsample was enriched for cases of substantial/major content change to facilitate human-model comparison. For the three-level primary content-change rating, agreement among the four raters was quadratic-weighted Cohen’s κ = 0.76 (range, 0.70–0.82; Krippendorff’s α = 0.77). The model reached quadratic-weighted κ = 0.75 against the mean rating across the four human raters and 0.69 against individual raters, and all ten model-rater and rater-rater comparisons reached substantial agreement or better (**Supplementary Fig. 2a,b)**. The model was within one ordinal level of the mean human rating on 98.7% of pairs; among the pairs with a mean human rating at either end of the scale (unchanged or major), 98.1% (362 of 369) were not assigned to the opposite end by the model (**Supplementary Fig. 2a**). Examples of major disagreement between the raters and the model are described in Supplementary Table 1. Across three repeated model runs, exact agreement on the primary content-change label was 84.2– 85.0% (unweighted Cohen’s κ = 0.73–0.75; quadratic-weighted κ = 0.79–0.80). On the major-versus-non-major classification, the model tracked rater conviction closely, classifying a revision as major in 5% of pairs where no rater did so and in 100% of pairs where all four did (**Supplementary Fig. 2c**). The model also reproduced the direction of the hedging shift. The excess of more-cautious over more-confident shifts was present in each of the four raters as well as in the model (**Supplementary Fig. 2d**). Across the 470 pairs with a unique human modal label, exact LLM–human agreement was 80.9%, while opposite-direction classifications occurred in only 0.9% of pairs (**Supplementary Fig. 2e**). Because the validation sample was deliberately enriched for difficult cases, its absolute shift frequency is not directly comparable with the full-corpus estimate; across all 514 assessable pairs, individual human shift rates ranged from 12.3% to 21.2%, compared with 27.0% for the LLM (**Supplementary Fig. 2f**).

Since pairwise extraction could inflate apparent claim stability by prompting the model to locate a corresponding claim in both versions, we tested this in a random 500-pair sample, processing each abstract independently, blind to its counterpart and to its version label. Independent and paired extraction selected the same or an overlapping formulation of the same primary claim in 96% of preprint and 95% of published abstracts; and the identical claim span in 80% and 78% (**Supplementary Fig. 3a**). A genuinely different claim was selected in 3.9% and 5.5% (**Supplementary Fig. 3a**). Classification agreement over major revision remained 96–100% across all but the lowest text-similarity stratum, where it was 84% (**Supplementary Fig. 3b**).

In a separate human validation of 100 pairs (200 abstracts), exact agreement between the model and individual human raters in primary-claim sentence selection ranged from 52.0% to 73.0%. Agreement increased to 76.9% when at least three raters selected the same sentence and to 90.0% when all four raters agreed (**Supplementary Fig. 3c**). Following this validation process, we applied the Sonnet model to the full corpus; the human labels served as an error check and were not used to tune the model or the codebook.

### Primary scientific claims usually persist but wording turns more cautious during preprint-to-publication transition

Most primary claims were preserved from preprint to peer-reviewed publication. Across all 72,644 abstract pairs, median Jaccard similarity was 0.838 (25th to 75th percentile, 0.654–0.949); 38.5% had similarity ≥0.90 and 5.9% had identical normalized word sets. The primary claim was unchanged in 39.8% of pairs and only minorly revised in 50.0%, leaving 10.2% with a major change in content, including 1.7% (n = 1,246) in which the primary claim was replaced, added or removed (**Fig. 1a**). When the certainty of language changed, it shifted toward caution more often than toward confidence. In the LLM-classified full corpus, hedging was unchanged in 85.6% of abstracts and could not be assessed in 1.8%, mostly because of such claim replacement. Among those with any hedging shift, twice as many primary claims became more cautious as more confident (8.4% versus 4.2% of all pairs; **Fig. 1b**). This shift toward caution increased with the extent of revision. Among abstracts with a major content change, hedging became more cautious in 38.5% of assessable claims and more confident in 19.8%, whereas abstracts with unchanged content rarely shifted in certainty (**Fig. 1c**). The excess of more-cautious over more-confident shifts was statistically significant (two-sided sign test on the 9,150 pairs with any hedging shift, P < 0.001).

**Figure 1:**
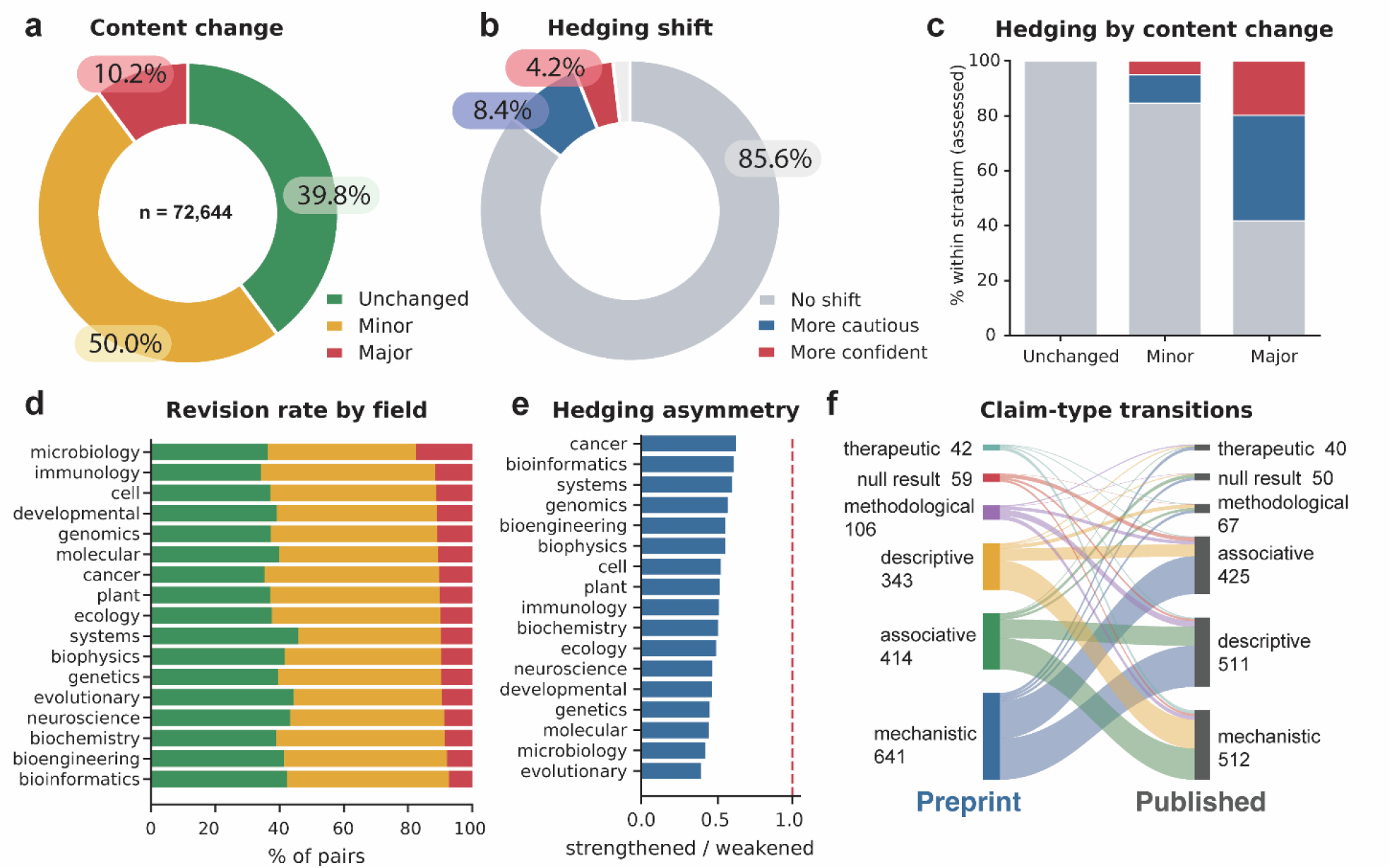
Patterns of preprint-to-publication changes in primary abstract claims. **a,** Content change of the primary claim across all 72,644 matched preprint-publication pairs. **b,** Hedging shift of primary claims. Hedging was non-assessable in 1.8% of pairs (n = 1,328), including 1,246 pairs (1.7%) with a replaced, added, or removed primary claim, and is shown in grey. **c,** Composition of hedging shift within each content-change stratum, expressed as a percentage of claims with an assessable hedging label. **d,** Content-change composition by field of bioRxiv subject category, ordered by the rate of substantial/major revision. Only fields with at least 1,500 pairs were included. **e,** Ratio of more confident (strengthened) to more cautious (weakened) hedging shifts by field; the dashed line marks parity. **f,** Alluvial diagram of primary claim-type transitions among the 1,605 out of the total 72,644 pairs with different valid claim types between preprint and publication, after excluding 1,246 pairs with a replaced, added, or removed primary claim and 343 pairs without valid claim-type assignments for both versions; among the 71,055 pairs with corresponding primary claims and valid claim types, 641 of 25,323 mechanistic, 414 of 11,946 associative, 343 of 13,478 descriptive, 106 of 16,435 methodological, 59 of 1,387 null-result and 42 of 2,486 therapeutic preprint claims changed type; band and bar widths are proportional to the number of pairs.

We next asked how the stability of scientific claims varied across the fields and claim types. Across the corpus, neuroscience was the largest field (18.6% of pairs), and the primary claims were predominantly mechanistic (35%) or methodological (23%; **Supplementary Fig. 4**). We observed substantial/major revision of the primary claim ranging from 7.2% of pairs in bioinformatics to 17.5% in microbiology (**Fig. 1d**). The shift toward caution was nonetheless stable in every field, with the ratio of more confident to more cautious primary claims below one in all 17 fields with at least 1,500 pairs (**Fig. 1e**, sign test p<0.001). Primary claim type was preserved in 97.7% of the 71,055 pairs with corresponding primary claims and valid claim-type assignments for both versions. Among the 2.3% (n = 1,605) that changed type, transitions occurred mostly between related categories, for example from mechanistic to associative or descriptive claims, rather than to a null result (**Fig. 1f**). In absolute numbers, mechanistic claims were most frequently reclassified, usually as descriptive or associative, and descriptive claims showed the largest net gain.

### Patterns of revision vary with claim type, preprint-to-publication intervals and journal impact

We next asked whether revision of the primary and secondary claims was correlated. Secondary claim 1 (S1) changed in 90% of pairs when the primary claim was substantially revised, compared with 34% when the primary claim was unchanged (chi-squared test, P < 0.01) (**Fig. 2a**). This suggests that substantial/major revision often affects the abstract claim structure in a coordinated manner, rather than isolated wording. Moreover, the stability of the primary claim varies with its type. The primary claim was substantially revised in only 5.4% of methodological claims, compared with 11.4% to 11.9% of descriptive, associative, and mechanistic claims (**Fig. 2b**). Within the same papers, the secondary claims were revised more often than the primary claim for every type, interestingly, with the largest difference observed for methodological claims (11.7% versus 5.4%; **Fig. 2b**).

**Figure 2:**
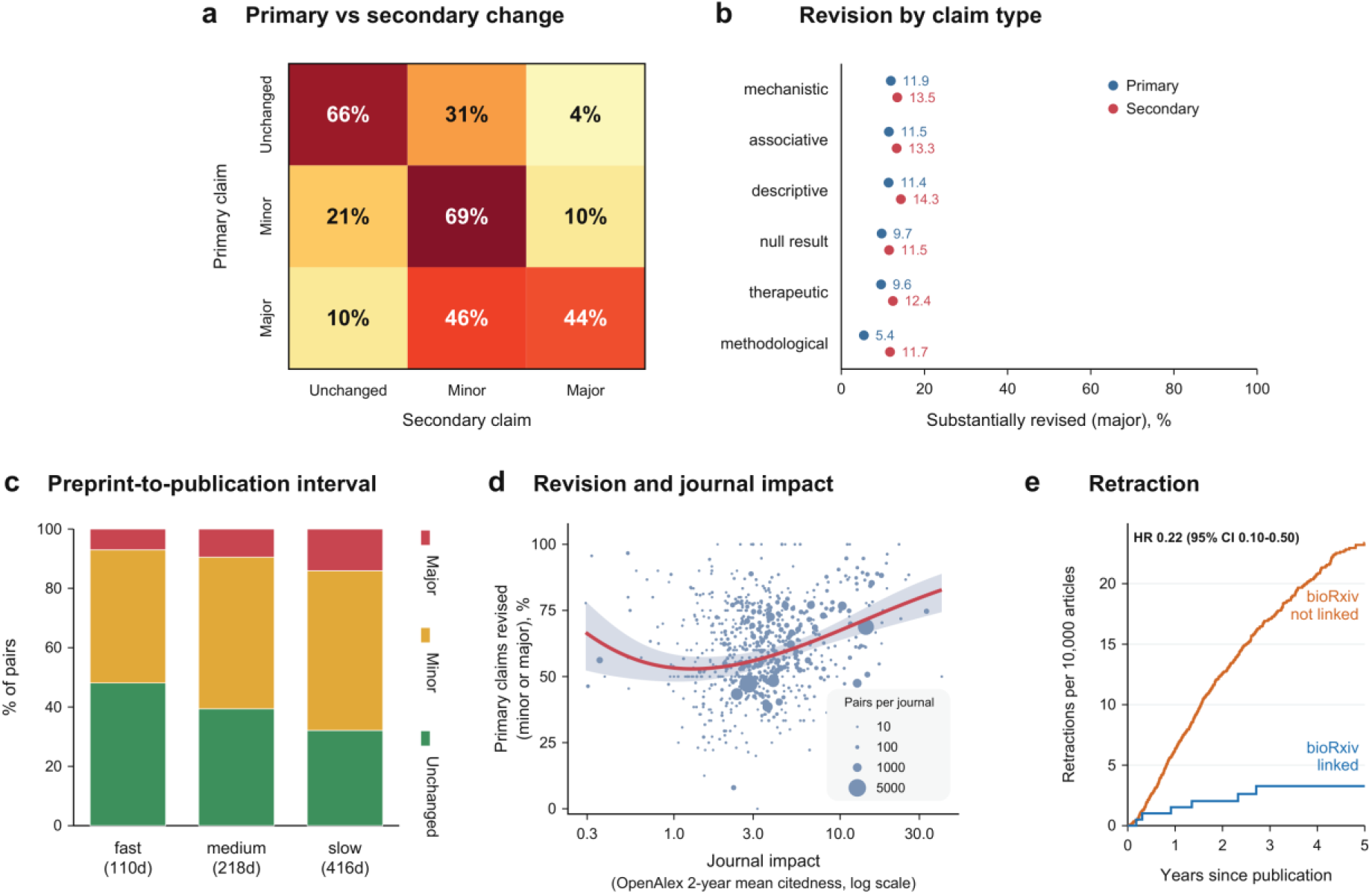
Factors associated with preprint-to-publication revision. **a**, Content change of secondary claim 1 (S1) (columns) as a function of the content change of the primary claim (rows); row values sum to 100%. **b**, Rate of substantial/major revision of the primary claim (blue) and of the pooled secondary claims (red) within each primary claim type, ordered by the primary rate. Claim type is defined by the primary claim. **c**, Content-change composition by tertile of preprint-to-publication interval, with the median number of days shown per tertile in parentheses. Red: major revision; yellow: minor revision; green: unchanged. d, Association between journal impact and any primary-claim revision among 59,012 pairs across 736 journals. Journal impact was measured using OpenAlex 2-year mean citedness. Each point represents one journal and is sized according to its number of pairs. The solid line shows the fitted probabilities from the selected grouped binomial FP2 model (powers −0.5 and 0); shading indicates the robust 95% confidence interval. e, Cumulative incidence of retraction for articles with (blue) or without (orange) a linked bioRxiv preprint. Hazard ratio (HR) and 95% confidence interval (CI) from a Cox model stratified by journal and publication year.

Substantial/major revision declined across years of preprint posting, from 17.0% of pairs posted in 2019 to 5.7% in 2024 (Supplementary **Fig. 5a**). The median preprint-to-publication interval also decreased from 666 to 160 days over this period (**Supplementary Fig. 5b**). We repeated the analysis among pairs published within 365 or 730 days, excluding posting dates for which the full corresponding publication window was not covered by the dataset. The association between posting year and substantial/major revision remained inverse in both analyses (**Supplementary Fig. 5c**). Because these analyses include only preprints that reached publication within the specified windows, manuscripts requiring substantial revisions and longer publication timelines may be underrepresented.

Furthermore, substantial/major revision increased with the preprint-to-publication interval, rising monotonically from 7.0% of pairs in the fastest tertile (median 110 days) to 14.1% in the slowest tertile (median 416 days; **Fig. 2c**, chi-squared test, P < 0.001). Continuous analysis confirmed a positive, nonlinear association, with predicted probabilities of substantial/major revision of 6.8% at 90 days, 12.3% at 365 days, and 17.5% at 730 days (**Supplementary Fig. 6**). Journal impact, measured using OpenAlex 2-year mean citedness, was nonlinearly associated with any primary-claim revision (grouped binomial fractional-polynomial regression, FP2 powers −0.5 and 0; P < 0.001 for nonlinearity; **Fig. 2d**). The fitted probability was lowest near a 2-year mean citedness of 1.3 and increased thereafter, from 53.8% at citedness 2 to 67.5% at citedness 10 (OR 1.78 comparing citedness 10 versus citedness 2, robust 95% CI 1.39–2.29).

Authors can post a preprint at any stage of a manuscript, from before first submission to after acceptance, so a version deposited shortly before publication may have peer review incorporated, which could inflate the apparent stability of claims. We therefore tested whether the preprints posted closest to publication, representing the plausible post-review deposits and with the shortest preprint-to-publication intervals, could account for the claim stability observed. We found that only 1.3% of preprints were posted within 30 days of their publication and 11.3% within 90 days, against a median interval of 218 days, and none were posted after publication (**Supplementary Fig. 7a)**. These short-interval pairs were more similar in wording to their published versions and were revised less often (**Supplementary Fig. 7b)**. However, excluding every pair posted within 90 days of publication barely changed the rate of substantial/major revision (from 10.2% to 10.7%; **Supplementary Fig. 7c**), suggesting that post-peer review deposits did not substantially contribute to the stability of primary scientific claims.

Additionally, an exploratory time-to-event analysis showed that preprinting was associated with a lower rate of later retraction. Among 399,928 original research articles published in 47 journals between 2021 and 2024, retraction occurred at 0.79 per 10,000 article-years among 19,692 articles linked to a bioRxiv preprint (6 events) and 5.20 among 380,236 without an identified preprint (715 events; hazard ratio 0.22, 95% confidence interval 0.10 to 0.50, P = 2.7 × 10^-4^, Cox model stratified by journal and publication year; **Fig. 2e**). The result was similar in every alternative model we tested, including one that corrects for the small number of events (Firth-penalized Cox model, hazard ratio 0.24, 95% confidence interval 0.10 to 0.48, P = 4.0 × 10^-6^) (**Supplementary Fig. 8**).

## Discussion

We compared the abstract claims of 72,644 bioRxiv preprints with their peer-reviewed publications. Nearly 90% of primary scientific claims were unchanged or only minorly revised, and substantial revision was uncommon. Where claims changed, the language shifted toward caution more often than toward confidence. Substantial/major revision was more frequently associated with longer preprint-to-publication intervals, and, in an exploratory time-to-event analysis, linkage to a bioRxiv preprint was associated with a lower rate of subsequent retraction.

These findings are consistent with smaller studies reporting that most preprint claims are retained through peer review^9,10,12,17^ and with a recent scoping review reaching the same conclusion across the health literature^11^. We extend this work to the full bioRxiv corpus. This corpus, by design, captures preprints that reached journal publication (previously estimated >70% of all bioRxiv preprints go on to be published in peer reviewed journals^2^); preprints without a matched publication were not represented in our analysis. A more important development is that we compare at the level of the scientific claim rather than textual similarity. This distinction matters, as two abstracts can be verbally different while preserving scientific claims, or verbally similar while changing scope, magnitude or even direction. The shift toward more cautious wording is consistent with the reduction in reported uncertainty during peer review^17^ and with evidence that reviewers or editors constrain overstatement in abstracts^18^. However, the observed differences cannot be attributed solely to journal peer review; they may also reflect feedback from readers following public posting or independent revisions by the authors. Changes prompted by community feedback would illustrate a potential benefit of early dissemination, whereas changes arising through journal review would reflect traditional editorial quality assurance. Our data cannot distinguish or quantify these contributions. The apparent decline in substantial revision over time may partly reflect the publication coverage of our dataset (January 2021 to February 2025). Recent preprints undergoing substantial revision may take longer to publish and therefore be less likely to enter our dataset before the cut-off. The fixed-window analyses ensure complete follow-up but remain conditional on publication within those windows, so the calendar-time findings should be interpreted as descriptive patterns.

Our exploratory retraction analysis revealed that linkage to a bioRxiv preprint was associated with a lower rate of subsequent retraction (hazard ratio 0.22, 95% confidence interval 0.10 to 0.50). This estimate rests on six events, at most 5.6 years of follow-up, and those preprints that ultimately reach journal publication. Although stratification by journal and publication year accounts for differences by venue and calendar time, residual confounding cannot be excluded; the direction is consistent with reports of comparable preprint quality^10,11^ and rare retraction of preprinted research^19^, and provides no support for the view that bioRxiv preprints are less reliable.

The shift toward more cautious language, even when central claims persist, can be important for how the readers perceive and use the findings. A published abstract with decreased language certainty or weakened effect type (e.g. from causal to associative language) may reflect peer reviewers’ comments against overstatement or revision efforts that demonstrate further complexity of the scientific observations. However, this line of information becomes available only upon publication, with a median of approximately seven months after a preprint is posted. Future studies will need to assess whether large language models can provide an equivalent calibration of claim strength at the time of posting, which could make this information available without delay directly on the preprint. Furthermore, because the corpus largely precedes the routine use of large language models in scientific writing, the trend from 2018 to 2024 also provides a reference point for assessing how the expansion of such tools modifies preprint-to-publication revisions^20,21^.

The positive correlation between the extent of revision and preprint-to-publication interval is consistent with a previous study based on linguistic distance, in which greater linguistic change was associated with a longer time to publication^16^. However, it should be noted that this cannot be interpreted as a causal effect of peer review, as longer intervals can be attributed to author choices over preprint deposit timing (i.e., before submission, during review or even after acceptance), review requirements, experimental nature, or field- or journal-specific practices.

We observed a non-linear association between journal impact and the frequency of primary claim revision. Above the fitted minimum at a 2-year mean citedness of approximately 1.3, revision frequency increased with citedness. Because this outcome includes minor revision, the rise may reflect more extensive editing of abstracts and more demanding review at more highly cited journals. Field, article type and time to publication also differ between journals and were not adjusted for, so this association should not be read as an effect of journal impact itself.

Post-review deposition of a preprint is a source of overestimation of claim stability. To address this consideration, we analyzed exclusively the first posted version of the preprint and performed sensitivity analyses through removing those with short preprint-to-publication intervals, which reduces this risk but does not remove it; residual cases cannot be fully excluded without confirming version status directly with the authors or comparing preprint posting dates with the submission and acceptance dates that some journals report.

Our study has several limitations. We analyzed abstracts rather than full texts, so changes confined to the methods, figures, or results were not captured. We compared the first preprint version with the published version. The observed changes combine author revision, inputs of reviewers and editors, and journal production. Claims were labeled by a single large language model. Despite the agreement between the model and the human experts, automated extraction remains imperfect^21^. In addition, our workflow required extraction of one primary claim and did not include a no-claim category; future studies could allow claims to be absent and classify the appearance or disappearance of a qualifying claim as a substantial revision. In this regard, we report the model, its version, and the prompts following current reporting standards^22,23^. We also did not separately validate the labels within individual fields or for the secondary claims, so the reported prevalences may carry field-specific or claim-specific measurement error. Because the corpus includes only published pairs, recently posted and slower-to-publish preprints are underrepresented, while preprints without a matched publication during the observation period are excluded. Preprints reaching publication may differ systematically from those without a matched publication, including in study quality, author resources, or scientific attention. This selection may accentuate the apparent decline in substantial revision over time and may overestimate claim stability across bioRxiv preprints generally. In addition, the corpus is restricted to bioRxiv and may not generalize to clinical, social science or other preprint literatures. Future studies could also complement this corpus-level analysis with detailed qualitative examination of smaller random samples, including version histories and individual revisions, and assess whether controversial or high-attention preprints exhibit distinct revision patterns.

On a general scale, preprinted manuscripts are not a fully representative sample of biomedical research. The decision to post a preprint, and the choice of which manuscript to post, likely depends on the authors and on how complete the work is at submission, so our estimates describe the preprinted literature rather than all biomedical manuscripts.

Taken together, the present results provide a quantitatively calibrated view of claim stability among bioRxiv preprints that subsequently reached journal publication. Among these preprints, the abstract claims usually persist, but published versions often experience the refinement of the language certainty and minor revisions of claims.^24^ Claim stability is not interchangeable with scientific correctness, which we did not assess directly, but it constitutes a necessary condition for using preprints as an early source of the biomedical knowledge.

## Methods

### Study design and sample

Using the bioRxiv API (/pubs/ endpoint) and PubMed metadata, all bioRxiv records posted between 2018 and 2025 that had a DOI corresponding to a peer reviewed original research article and that were published between January 2021 and February 2025 were retrieved. We selected 2018 as the starting year because new bioRxiv preprints increased substantially from 2018 (with more than 20,000 preprints per year^25^), providing sufficient annual sample sizes for temporal analyses. No statistical methods were used to predetermine sample size; the analysis includes every preprint-publication pair meeting the eligibility criteria.

Pairs were included in the analysis if both abstracts were in English and contained at least 100 characters. For manuscripts with multiple preprint versions, only the first version was included. The final corpus consisted of 72,644 matched preprint-peer reviewed publication abstract pairs, which cover 3,149 journals and 27 bioRxiv subject categories (the 17 with at least 1,500 pairs are shown in the field analyses). Of 74,081 pairs passing the abstract-length filter (of 74,098 matched pairs), 72,644 (98.1%) received complete Sonnet labels; 1,437 (1.9%) failed because of transient Sonnet API errors during batch processing. These pairs were not rerun after the extraction batch was closed and the analytical dataset frozen; this was a technical exclusion rather than an assessment of abstract suitability. For retraction rate analysis, an article-level cohort was assembled for the same journals over the same period (see statistical analysis); retraction data were obtained from the Retraction Watch database distributed by Crossref.

### Claim extraction and classification

For each preprint–publication abstract pair, the Claude Sonnet 4.6 model (Anthropic) was used to extract (1) one primary claim and up to two secondary claims, (2) a claim type (mechanistic, associative, descriptive, methodological, therapeutic, or null result), and (3) a hedging level indicating the tier of language certainty. When two secondary claims were present, they were ranked by mechanistic relevance, then by moderator or scope content over applied implications, followed by order of appearance and sentence-length proximity to the primary claim’s length. These are referred to as secondary claim 1 (S1) and secondary claim 2 (S2).

With a second prompt, the model was used to compare the claim of each preprint to that of the published counterpart. Quantitative aspects of the claims included:

1. Content change, which was categorized into 3 levels: unchanged, minor revision, or substantial/major revision. A claim was labelled as unchanged when the published version was identical to or trivially paraphrased from the preprint claim, without changes in direction, scope, magnitude, effect type or certainty. A claim was labelled as minor revision when the change was non-substantive, including wording-only paraphrases or hedging shifts that did not alter the scientific content. A claim was labelled as major revision in the presence of the following changes: direction flips, scope changes (entity, population, species, setting), magnitude shifts ≥ 20 % on a comparable estimator, effect-type transitions (associative, predictive, explanatory, causal regulation, causal necessity), or outright claim replacement. Transitions between presence and absence of an effect were treated as substantial/major revision. Rhetorical modifications, including language/phrases of novelty or potential importance, were not considered as scope changes. This categorization was modified from a prior study^26^ to fit the LLM prompt.
2. Hedging shift, which was categorized into 3 levels: more cautious, unchanged, more confident. A primary claim that was replaced, added, or removed was classified as a major revision and assigned “not applicable” (NA) for hedging. These pairs did not contribute to the more-cautious or more-confident categories. Certainty was assessed using verb strength and causal language. The verbs with high certainty were “demonstrates, shows, establishes, proves, reveals, causes, is required for, is necessary for, is sufficient for”; verbs with moderate certainty were “suggests, indicates, supports, is consistent with, provides evidence that, implies”; verbs with low certainty were “may, might, could, can, potentially, possibly, appears to, seems to, speculate, hypothesize”. Causal language of the claims was ranked from weaker to stronger effect type classes: associative, predictive, explanatory, causal regulation, and causal necessity, following established ordered scales of causal-claim strength in which movement up the scale denotes a stronger claim.^27,28^. Movement upward in this rank was classified as more confident, whereas downward as more cautious. Any hedging shift makes content change labelled at least minor change, even when the content of scientific claim was preserved.

All calls used Claude Sonnet 4.6 (model identifier claude-sonnet-4-6) at temperature 0 with a 1,200-token output limit, returning structured JSON under the locked v7.1 codebook (locked 25 April 2026).

To assess whether paired preprint-publication presentation to LLM influenced claim extraction, we repeated extraction with each version processed separately (without preprint/publication labels or access to its counterpart) in a prespecified 500-pair sample using the same locked codebook, and compared the results with the pipeline. Selections were scored as identical when the spans matched after normalisation, or differed only by an implication or elaboration clause that the codebook’s scope rule excludes from the claim; other overlaps were scored as partial. Human-model extraction agreement was evaluated separately in 100 randomly sampled pairs (200 abstracts). Four raters viewed both abstracts without titles or access to the model selections and independently selected the sentence containing the primary claim. Agreement was defined as selection of the same sentence and summarized for each individual rater and according to the level of agreement among the four raters. Confidence intervals were obtained by cluster bootstrap over preprint-publication pairs.

To establish the reliability of the content-change and hedging labels, four raters independently labelled a stratified subsample of 550 abstract pairs using the pre-registered codebook (v7.1), blind to the model output. Raters were shown the primary claims of the preprint and the published version, with both abstracts available for reference, and rated the content change (unchanged, minor or major) and hedging shift (more cautious, unchanged or more confident) of each pair. The validation sample comprised 260 representative pairs, 80 additional field-stratified random pairs, 40 disagreement-enriched pairs, and 170 pairs enriched for model-classified major revisions. For content change, the aggregate human rating was calculated as the mean ordinal rating of the four raters (unchanged = 0, minor = 1, major = 2), binned at 0.5 and 1.5, with exact ties assigned to the more severe category. For hedging shift, pairs whose primary claim was replaced outright were excluded, leaving 514 assessable pairs. Comparisons with individual human raters used all 514 pairs, whereas comparisons with the human modal label used the 470 pairs with a unique mode; the remaining 44 pairs were retained for individual-rater comparisons. Agreement on content change was quantified with quadratic-weighted Cohen’s κ and Krippendorff’s α. For hedging shift, we report exact agreement and opposite-direction classifications between the model and the aggregate human label. Confidence intervals are 1,000-iteration bootstrap percentiles. Three replicate Sonnet runs at temperature 0 were compared for reproducibility of the primary content-change label; the resulting agreement statistics are reported in the Results. The single Sonnet scheme was then applied to the full corpus, apart from 3,010 pairs labelled in an earlier pilot batch that retain three-run majority labels; these differ from the single-pass labels in the boundary between unchanged and minor revision but not in the rate of substantial/major revision, and excluding them shifts the overall distribution by less than 0.5 percentage points. All values were computed in Python or R.

### Statistical analyses

Claim-change prevalence was estimated as the proportions of primary claims categorized as unchanged, minor revision, or major revision, and 95 % Wilson confidence intervals of each category. Estimates were stratified by field, claim type, journal impact, calendar year of preprint posting (2018–2025), and preprint-to-publication interval tertiles (fastest to slowest, calculated as interval from preprint to publication). Journal impact was the 2-year mean citedness from OpenAlex (matched by journal name; available for 736 journals covering 59,012 pairs), which is different from the Journal Impact Factor. Preprint-to-publication interval was defined as the interval in days from the first-version preprint posting to journal publication, split into equal-sized tertiles (medians 110, 218, and 416 days). The relationship in content change between primary and secondary claim was examined using a chi-squared test.

Hedging shifts between preprints and publications were analyzed for the frequency and direction (more cautious vs. more confident). A Wilcoxon signed-rank test on paired hedging scores and an exact two-sided sign test were used to evaluate the asymmetry between more cautious shifts and more confident shift in the claims with any hedging shifts. Ratios of more-confident to more-cautious shifts were examined across fields and claim types. Claim-type transitions were counted and visualized using an alluvial diagram after excluding pairs with replaced, added, or removed primary claims and pairs without valid claim-type assignments for both versions.

The association between substantial/major revision and year of preprint posting was assessed using logistic regression, with posting year entered as a continuous variable. Sensitivity analyses were restricted to pairs published within 365 or 730 days and to posting dates for which the full corresponding publication window fell within the dataset’s publication coverage from 1 January 2021 through 28 February 2025. Eligible posting dates ranged from 1 January 2021 through 29 February 2024 for the 365-day analysis and from 1 January 2021 through 1 March 2023 for the 730-day analysis. These analyses remained restricted to preprints that reached journal publication within the specified windows. The tertile-based comparison of preprint-to-publication intervals was used as a descriptive analysis. For continuous inference, substantial/major revision was modeled using fractional-polynomial logistic regression^29^ with the standard power set p = {−2, −1, −0.5, 0, 0.5, 1, 2, 3} and a maximum degree of FP2, and closed testing at α = 0.05; the selected model had powers (1,1), corresponding to the terms x and x * log(x).

At the journal level, the numbers of pairs with any primary-claim revision (minor or major) and with an unchanged primary claim were modeled as grouped binomial outcomes, so each journal contributed according to its number of pairs. OpenAlex 2-year mean citedness was entered continuously, and its functional form was selected using fractional polynomials with the standard power set, a maximum degree of FP2, and closed testing at α = 0.05. The selected model had powers (-0.5,0), corresponding to the terms x^-0.5^ and log(x). Confidence intervals used HC1 heteroskedasticity-robust covariance estimates, treating journals as independent units.

To compare retraction rates between publications with and without a prior preprint, we built an article-level cohort for the 47 journals with unambiguous ISSN matches to our bioRxiv-linked publications. All Europe PMC records first published between 2018 and 2024 were retrieved and non-research publication types and records with missing or duplicated DOI/date excluded (29,938 and 1,970, respectively, of 641,907 records, 5.0%), leaving 609,998 articles. Exposure was linkage of the article DOI to a bioRxiv preprint; retractions and their dates came from the Retraction Watch database distributed by Crossref (snapshot 19 August 2026), which also set the censoring date. Because linked publications begin in 2021, the primary analysis covered 2021 to 2024 (n = 399,928; 721 retractions). Hazard ratios came from Cox models stratified by journal and publication year, with proportional hazards checked by Schoenfeld residuals (P = 0.33). Sensitivity analyses used a Firth-penalized model, the full 2018 to 2024 window, and an outcome including expressions of concern. The analysis was performed using R (4.3.3) packages survival and coxphf.

## Pre-registration

This work has been pre-registered on June 11^th^, 2026: https://www.researchhub.com/proposal/32332/tracking-claim-changes-from-preprint-to-publication-across-biomedical-studies-using-large-language-models

## Data and code availability

All original data are based on publicly available preprints and journal articles. The analysis code and LLM-derived full dataset are available on GitHub: https://github.com/rustlab1/PreprintPaperTracker and searchable on this website: https://rustlab1.github.io/PreprintPaperTracker/. The code and data have also been archived on Zenodo with a persistent DOI (https://doi.org/10.5281/zenodo.22664995).

## Acknowledgements

This work was supported by a USC Dean’s Pilot Award (to R.R.) and the ResearchHub Foundation (to H.Y. and R.R.). The funders had no role in study design, data collection and analysis, decision to publish or preparation of the manuscript.

## Author contributions

R.R. and H.Y. designed the study. R.R. built the pipeline and performed the analysis. H.Y., W.A., P.M.F. and R.R. validated the labels. R.R. and H.Y. wrote the manuscript. All authors revised and approved the manuscript.

## Competing interests

The authors declare no competing interests.

## Notes

https://rustlab1.github.io/PreprintPaperTracker

### Competing Interest Statement

The authors have declared no competing interest.

### Summary of Updates

Updated the model-human validation comparison and agreement statistics using the same primary-claim statements. Added examples of major disagreement in Supplementary Table 1 and revised Supplementary Figure 2. Added fixed-window analyses of calendar-time trends and clarified their interpretation. Revised Figure 2, figure legends, and discussion of journal impact and retraction findings. Updated the supplementary material.

