## Supplementary Information and Supplementary Table 1 for "Tracking claim changes from preprint to publication across 72,644 biomedical studies using large language models"

### SUPPLEMENTARY FIGURES

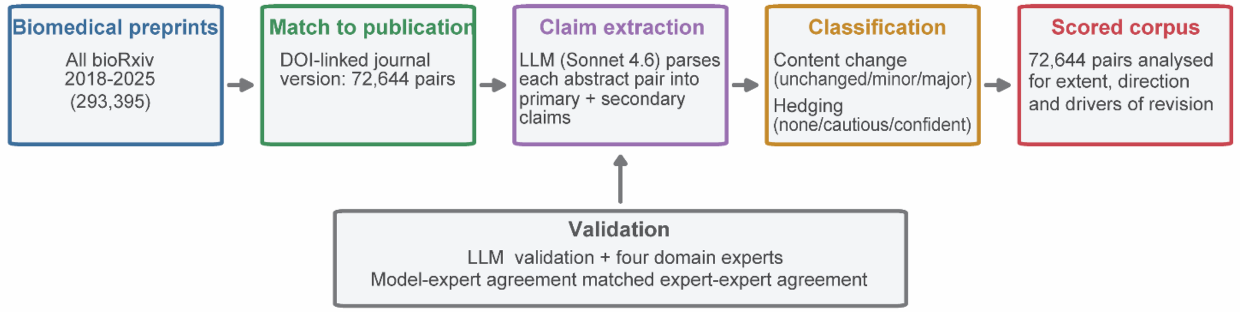


**Supplementary Figure 1: Study workflow. Corpus construction, claim extraction, classification and validation.** All bioRxiv preprints posted between 2018 and 2025 were screened and DOI-matched to peer-reviewed publications, yielding 72,644 preprint-publication pairs. Claude Sonnet 4.6 extracted one primary and up to two secondary claims from each abstract pair and classified primary-claim content change, hedging shift, and claim type under the locked codebook. Labels were validated against four independent human raters before the pipeline was applied to the full corpus.

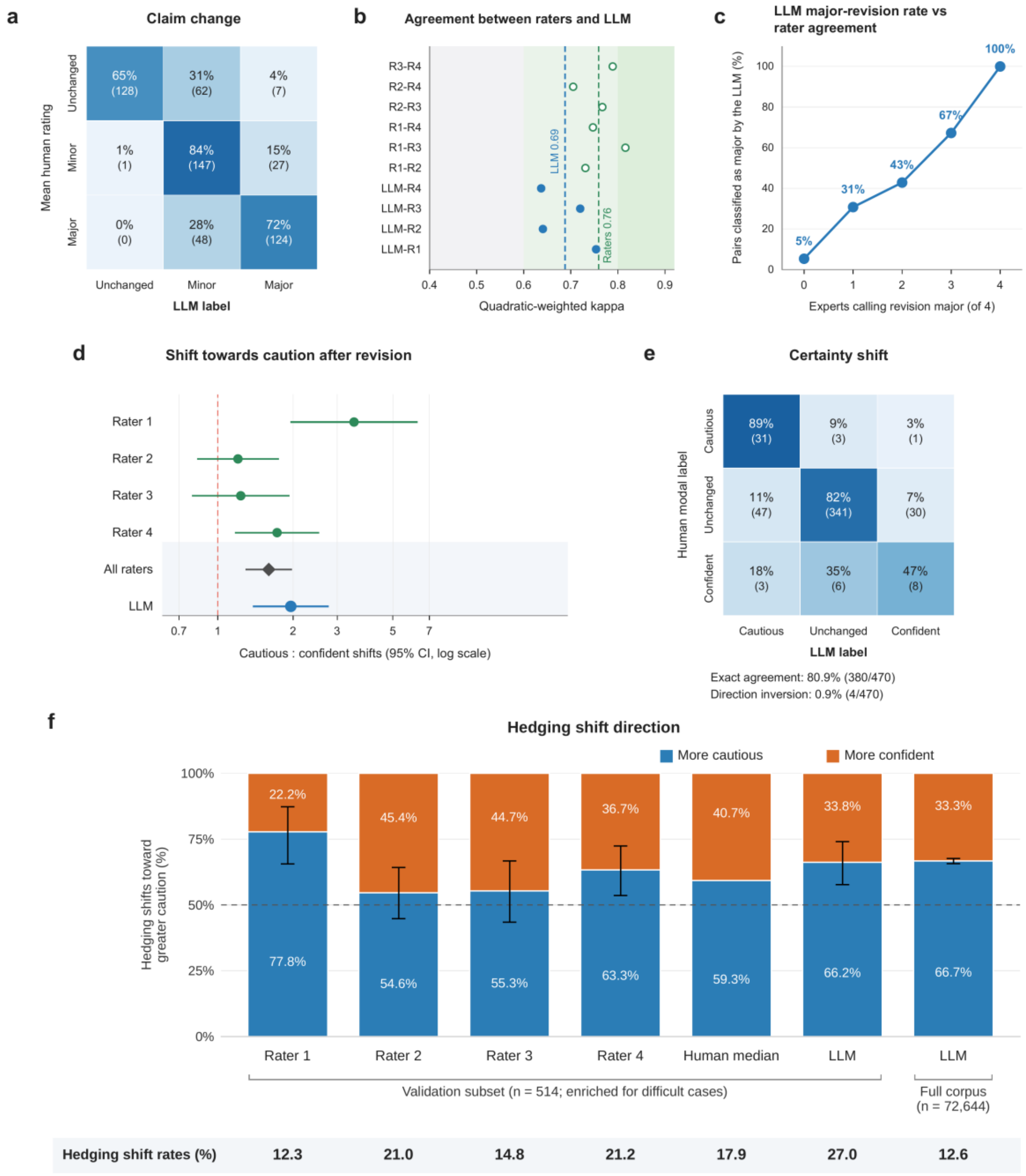


**Supplementary Figure 2: Reliability of the model labels against four independent raters.** **a**, Concordance between the model and the mean rating across four human raters for the three-level primary content-change classification (n = 544 pairs; model labels obtained on the claim statements shown to the raters); cells show row percentages with counts in parentheses. **b**, Quadratic-weighted Cohen's kappa for each model-rater comparison (filled circles) and rater-rater comparison (open circles); dashed lines show the corresponding mean values, and shading denotes the Landis–Koch interpretation bands. **c**, Percentage of pairs classified as a major revision by the model according to the number of human raters independently assigning a major revision. **d**, Ratio of more-cautious to more-confident hedging shifts for each rater, the four raters pooled, and the model (n = 514 assessable pairs); points show ratios, error bars show 95% Wilson confidence intervals, and the dashed line marks parity. **e**, Concordance between the model and the human modal hedging-shift label among 470 pairs with a unique human mode. Exact agreement comprised the diagonal cells (31 + 341 + 8 = 380), whereas opposite-direction classifications comprised the opposing corner cells (1 + 3 = 4). **f**, Direction of hedging shifts for each human rater, the median of the four raters and the model in the validation subset, and for the model in the full corpus. Stacked bars show the proportions of more-cautious and more-confident classifications among pairs identified as having a hedging shift; the row below the bars gives the hedging-shift rate (percentage of pairs with any shift). Rater and model estimates in the validation subset use all 514 assessable pairs; the human median bar shows the median of the four rater values and has no error bar. The validation subset was enriched for difficult cases, so its shift rates are not directly comparable with the full corpus. Error bars show exact 95% confidence intervals for the proportion classified as more cautious, and the dashed line marks equal frequencies.

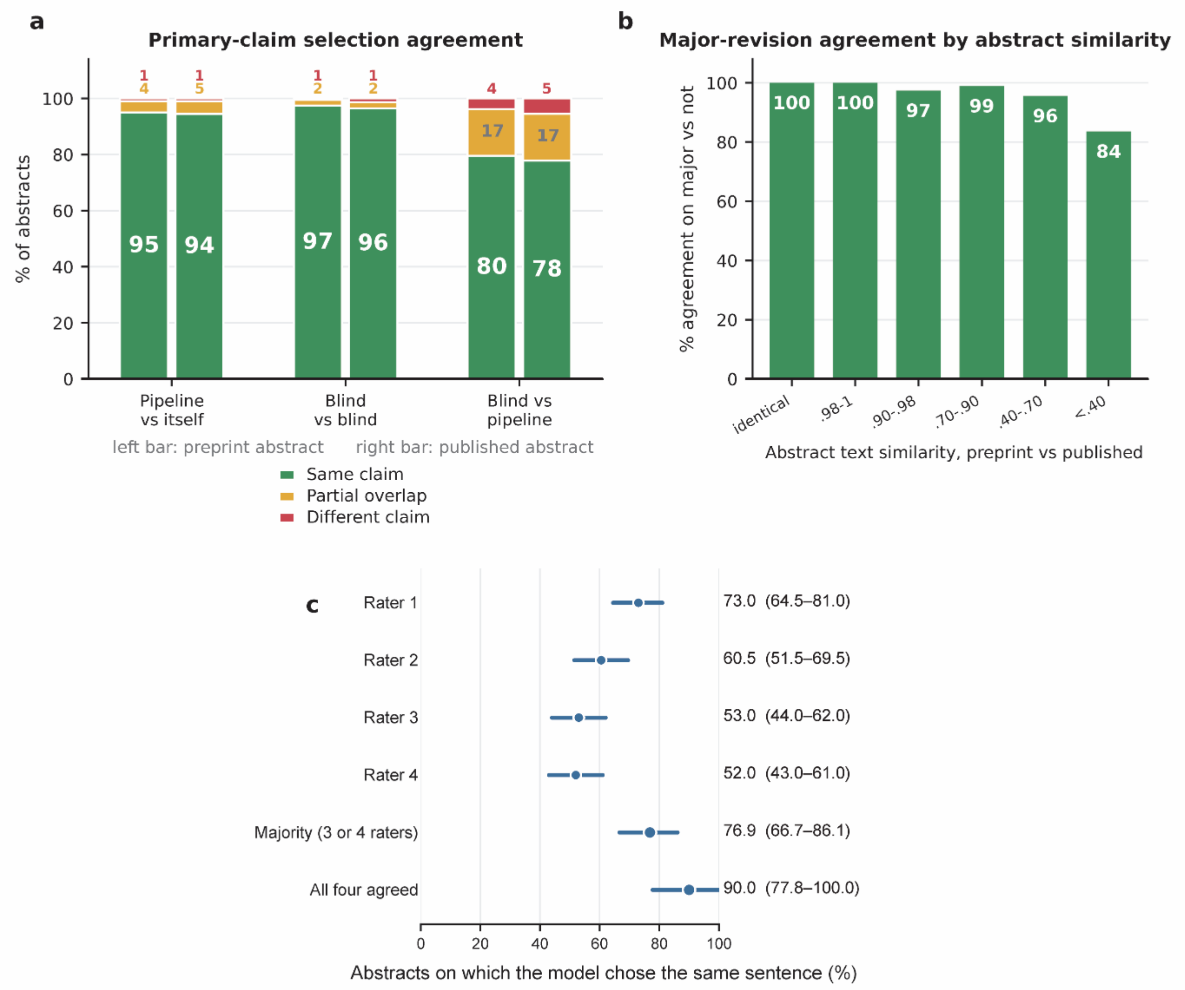


**Supplementary Figure 3: Blind independent claim-extraction sensitivity analysis.** Preprint and publication abstracts were processed independently, without indicating version status or providing the counterpart abstract. We refer to this as “blind” extraction, in contrast to “pipeline” extraction, in which the model receives both abstracts together. **a**, Agreement in primary-claim selection between repeated pipeline extractions (n = 598 pairs), repeated blind extractions (n = 593 pairs), and blind versus pipeline extraction in the prespecified 500-pair sample (n = 494 preprints and 492 publications with valid outputs). Left and right bars represent preprint and published abstracts, respectively. Colors distinguish the same claim, partial overlap, and a different claim. Consistent with the codebook's scope rule, partial-overlap cases were counted as the same claim when one extraction was a nearly complete trimmed span of the other. **b**, Agreement between the pipeline and blind procedures for the major-versus-non-major classification across strata of abstract text similarity. **c**, Exact agreement between model and human primary-claim sentence selection in 100 randomly sampled pairs (200 abstracts). Points show agreement for each individual rater and according to the level of agreement among the four raters. Error bars show cluster-bootstrap 95% confidence intervals over preprint-publication pairs.

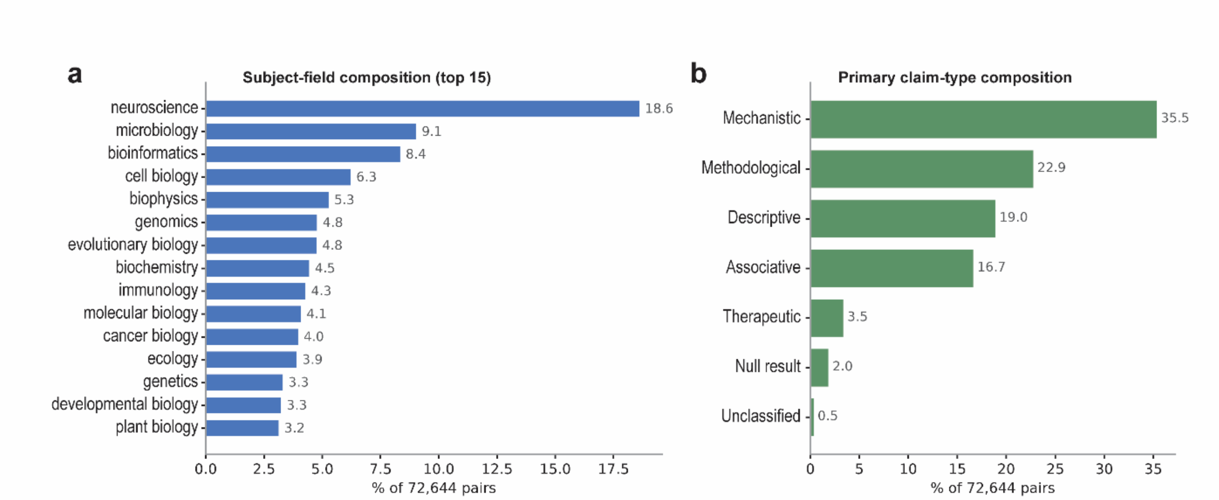


**Supplementary Figure 4: Composition of the corpus. a**, Distribution of the 72,644 preprint-publication pairs across the 15 most frequent bioRxiv subject categories. **b**, Distribution of primary claim types. Unclassified denotes the 343 pairs without a valid assignment to one of the six prespecified claim types. Percentages are calculated across all 72,644 pairs.

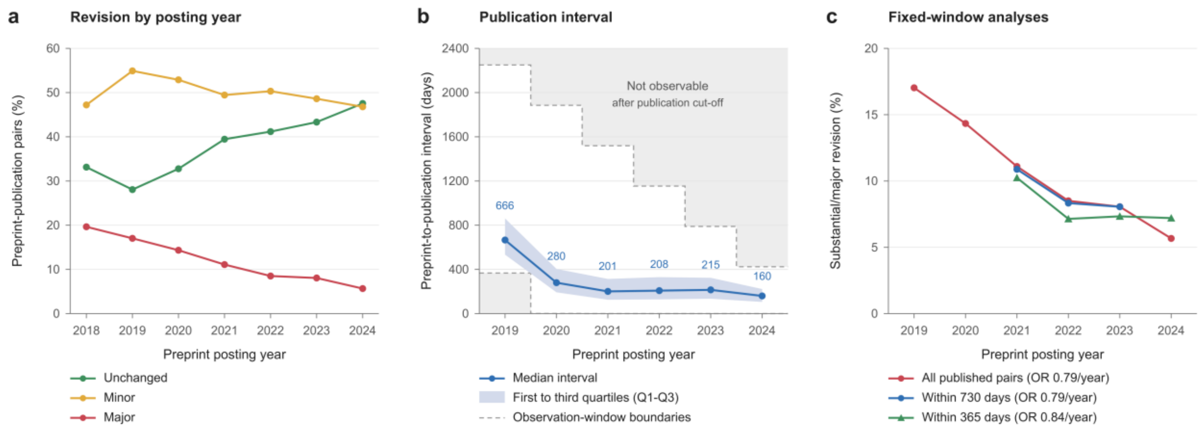


**Supplementary Figure 5: Calendar-time sensitivity analyses. a**, Content-change rates of primary claims by year of preprint posting. b, Median preprint-to-publication interval (points) and 25th to 75th percentile (shading) by year of preprint posting. Dashed lines mark the observable range given the publication coverage of the dataset (1 January 2021 to 28 February 2025); grey regions were not observable. Intervals above the upper line correspond to publication after the cut-off. For preprints posted in 2019, intervals shorter than 367 days correspond to publication before coverage began; this floor is 550 days for preprints posted mid-2019 and falls to 1 day by the end of 2020. c, Percentage of primary claims with a major revision by posting year for all pairs and for pairs published within 730 or 365 days. Fixed-window analyses were restricted to posting dates with complete corresponding follow-up. Odds ratios per year from logistic regression are shown in the key.

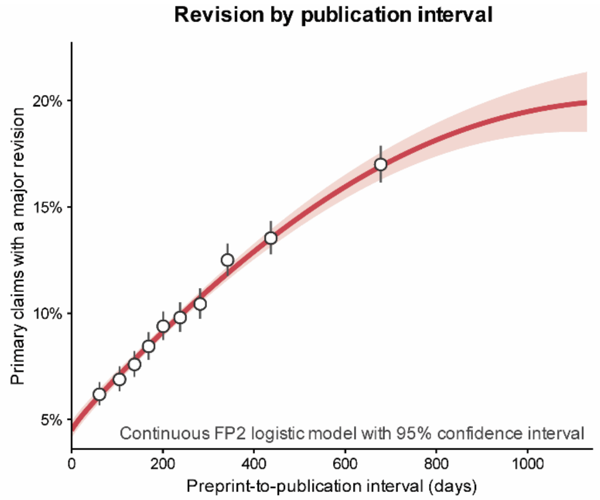


**Supplementary Figure 6: Continuous association between preprint-to-publication interval and substantial/major revision.** Circles show observed major-revision rates within interval bins, with 95% confidence intervals. The solid curve and shading show fitted probabilities and the 95% confidence interval from the selected fractional-polynomial logistic regression model (FP2 powers 1 and 1, i.e. x and xlog(x)).

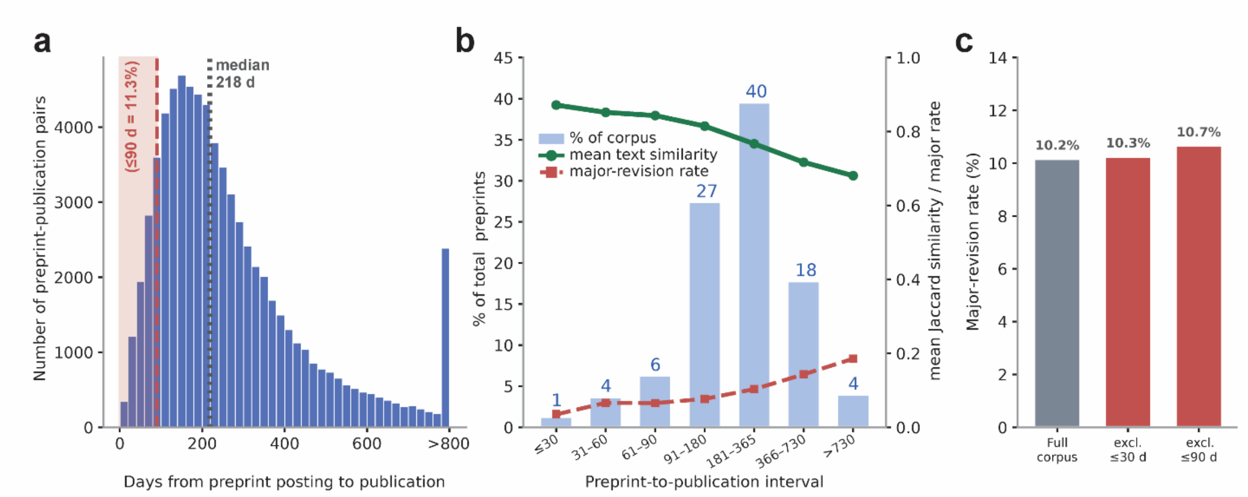


**Supplementary Figure 7: Short preprint-to-publication intervals do not explain primary-claim stability. a**, Distribution of the interval in days between preprint posting and journal publication; dotted line marks the median of 218 days; the shaded region marks pairs posted within 90 days of publication (11.3%). Intervals longer than 800 days were grouped into the final histogram bin, labeled “>800 days”. **b**, Corpus composition (bars and left axis), mean Jaccard similarity (green line, right axis), and major revision rate (red dashed line) by interval bucket of preprint-to-publication interval. **c**, Major revision rate for the full corpus (left bar) and after excluding pairs posted within 30 (middle bar) or 90 days of publication (right bar).

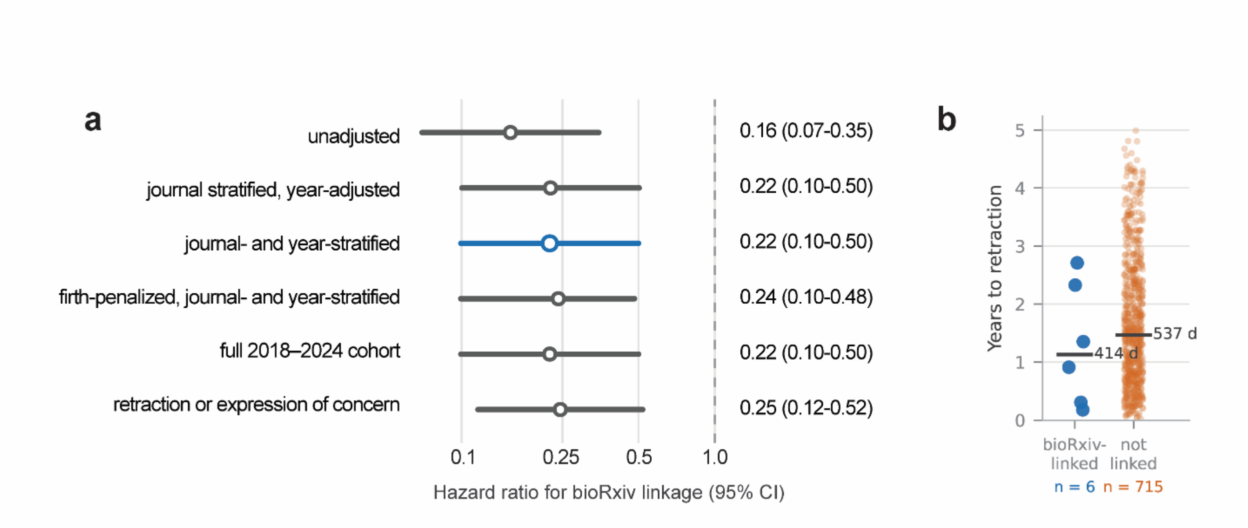


**Supplementary Figure 8: Sensitivity analyses of the association between bioRxiv linkage and retraction.** The analysis included 399,928 original research articles published during 2021 to 2024 in 47 journals, including 19,692 articles linked to a bioRxiv preprint and 380,236 without an identified bioRxiv preprint. **a**, Hazard ratio for bioRxiv linkage across model specifications. Points show estimates, horizontal lines show 95% confidence intervals, and the dashed line at 1.0 marks no association. The highlighted primary model is a Cox proportional-hazards model stratified by journal and publication year. Unadjusted is the crude model; journal stratified, year-adjusted includes publication year as a covariate within journal strata; journal- and year-stratified is the primary model; Firth-penalized, journal- and year-stratified applies a Firth penalty within the same strata to correct for the small number of events in the linked group; the remaining sensitivity analyses extend the period to 2018 to 2024 or include expressions of concern in the outcome. The stratified analysis comprised 187 journal-by-year strata, of which 50 contained both a retraction and a bioRxiv-linked article; all 721 events were retained. Because the coxphf package does not support strata, the Firth model used a one-parameter stratified penalised partial likelihood implemented for this analysis, which reproduced coxphf without strata and ordinary stratified Cox with Breslow ties to within $1 \times 10^{-7}$ on the log hazard ratio. **b**, Time from publication to retraction for each retracted article. Horizontal bars show group medians. Six events occurred in the bioRxiv-linked group and 715 in the unlinked group; the panel is descriptive.

**Supplementary Table 1: Validation pairs with major disagreement between the raters and the model.**

| **Pair 1. Genetics; preprint posted 2021; published in Human Molecular Genetics; bioRxiv DOI 10.1101/2021.10.01.462704** | |
|---|---|
| **Ratings** | **Raters 1–4:** unchanged, unchanged, unchanged, unchanged; mean 0.0, aggregate unchanged.<br>**Model:** major; hedging weakened. |
| **Claims shown to raters and model** | **Preprint:** Loss of α-actinin-3 in aged mdx dystrophic mice reduces pathological fibre branching and confers protection from eccentric contraction-induced force deficit through increased SERCA1 pump density and altered oxidative metabolism.<br>**Published:** Loss of α-actinin-3 in aged mdx mice reduces fiber branching complexity and protects fast-twitch EDL muscles from eccentric contraction-induced force deficit. |
| **Summary of model rationale** | The published claim drops the mechanism stated in the preprint (increased SERCA1 pump density and altered oxidative metabolism); classified as a scope and effect-type change (major), with hedging weakened. |

| **Pair 2. Biophysics; preprint posted 2021; published in Nature Communications; bioRxiv DOI 10.1101/2021.01.05.424557** | |
|---|---|
| **Ratings** | **Raters 1–4:** minor, unchanged, unchanged, unchanged; mean 0.25, aggregate unchanged.<br>**Model:** major; hedging weakened. |
| **Claims shown to raters and model** | **Preprint:** Positive allosteric modulators are required to stabilize the active state of mGlu2 receptors, as glutamate alone only partially stabilizes the extracellular domains while full activation occurs only in the presence of a PAM or Gi protein.<br>**Published:** Positive allosteric modulators enhance agonist efficacy by stabilizing the active state of mGlu2 receptors, increasing the residence time in the active conformation. |
| **Summary of model rationale** | The preprint states that positive allosteric modulators are required to stabilize the active state (causal necessity), whereas the published claim states that they enhance agonist efficacy (causal regulation); classified as an effect-type change (major), with hedging weakened. |

| **Pair 3. Evolutionary biology; preprint posted 2023; published in Peer Community Journal; bioRxiv DOI 10.1101/2023.07.06.547897** | |
|---|---|
| **Ratings** | **Raters 1–4:** unchanged, unchanged, unchanged, unchanged; mean 0.0, aggregate unchanged.<br>**Model:** major; hedging not assessable (claim replaced). |
| **Claims shown to raters and model** | **Preprint:** Aphid, an approximate maximum-likelihood method, successfully distinguishes gene flow from incomplete lineage sorting by analyzing branch lengths of three-species gene trees and quantifies their relative contributions to phylogenetic conflict.<br>**Published:** A substantial fraction of the human/chimpanzee/gorilla phylogenetic conflict is due to ancient gene flow. |
| **Summary of model rationale** | The preprint claim describes a method (Aphid) that distinguishes gene flow from incomplete lineage sorting, whereas the published claim states an empirical finding on human, chimpanzee and gorilla phylogenetic conflict; classified as replacement of the primary claim (major). |

| **Pair 4. Genomics; preprint posted 2020; published in PLOS ONE; bioRxiv DOI 10.1101/2020.09.15.297663** | |
|---|---|
| **Ratings** | **Raters 1–4:** minor, unchanged, unchanged, unchanged; mean 0.25, aggregate unchanged.<br>**Model:** major; hedging weakened. |
| **Claims shown to raters and model** | **Preprint:** P. agglomerans KM1 contains 13 antibiotic resistance genes, virulence factors including a type VI secretion system and hemolysin, and genes conferring resistance to environmental stresses, which collectively drive its pathogenic potential<br>**Published:** The P. agglomerans KM1 genome contains 13 antibiotic resistance genes, a type VI secretion system, hemolysin, filamentous hemagglutinin, iron uptake genes, and 6 CRISPR loci, and the isolate exerts strong immunostimulatory properties on macrophages through TLR4/TLR1/2 and NF-κB/MAPK-dependent pathways. |
| **Summary of model rationale** | The published claim adds filamentous hemagglutinin, iron uptake genes, CRISPR loci and immunostimulatory effects on macrophages, and drops the statement that these factors drive pathogenic potential; classified as a scope and effect-type change (major), with hedging weakened. |

| **Pair 5. Neuroscience; preprint posted 2021; published in Human Brain Mapping; bioRxiv DOI 10.1101/2021.01.30.428936** | |
|---|---|
| **Ratings** | **Raters 1–4:** unchanged, unchanged, unchanged, unchanged; mean 0.0, aggregate unchanged.<br>**Model:** major; hedging not assessable (claim replaced). |
| **Claims shown to raters and model** | **Preprint:** Body identity is decoded consistently across viewpoint in the fusiform body area, right anterior temporal cortex, middle frontal gyrus, and right insula, demonstrating that these regions extract high-level identity information similar to their function for faces.<br>**Published:** The brain encodes face and body identity in a partially separated manner, with distributed networks including fusiform body area, anterior temporal cortex, and early visual regions showing viewpoint-invariant identity decoding, while also maintaining a shared abstract person identity representation across faces and bodies. |
| **Summary of model rationale** | The preprint claim concerns viewpoint-invariant decoding of body identity in four regions, whereas the published claim describes partially separated coding of face and body identity, including early visual regions, and a shared person-identity representation; classified as replacement of the primary claim (major). |

| **Pair 6. Genomics; preprint posted 2023; published in Nature Communications; bioRxiv DOI 10.1101/2023.06.29.547066** | |
|---|---|
| **Ratings** | **Raters 1–4:** unchanged, unchanged, unchanged, minor; mean 0.25, aggregate unchanged.<br>**Model:** major; hedging not assessable (claim replaced). |
| **Claims shown to raters and model** | **Preprint:** Expression- and splicing-mediating variants are more than three times as frequent in testis compared to epididymis and vas deferens, highlighting the transcriptional complexity of testis.<br>**Published:** Regulatory loci identified through molecular QTL analysis in reproductive tissues reveal genes (WDR19, SPATA16, KCTD19, ZDHHC1) and molecular phenotypes associated with quantitative variation in male fertility in cattle. |
| **Summary of model rationale** | The two claim statements are different findings. |

| **Pair 7. Systems biology; preprint posted 2020; published in Scientific Reports; bioRxiv DOI 10.1101/2020.02.04.933713** | |
|---|---|
| **Ratings** | **Raters 1–4:** unchanged, minor, unchanged, unchanged; mean 0.25, aggregate unchanged.<br>**Model:** major; hedging unchanged. |
| **Claims shown to raters and model** | **Preprint:** Reinterpreting the interaction between PTB and nPTB genes is necessary to capture quantitative gene interaction dynamics and correctly predict the outcome of overexpression and knockdown experiments in direct neuronal reprogramming.<br>**Published:** nPTB feeds back into the direct neural conversion network via PTB to control gene interaction dynamics and predict outcomes of overexpression and knockdown experiments in neuronal reprogramming. |
| **Summary of model rationale** | The preprint claim states that reinterpreting the PTB–nPTB interaction is necessary to predict reprogramming outcomes, whereas the published claim states that nPTB feeds back into the network via PTB to control gene interaction dynamics; classified as a change in entity and effect type (major), with hedging unchanged. |
